# To buffer or to be labile? A framework to disentangle demographic patterns and evolutionary processes

**DOI:** 10.1101/2021.04.12.439165

**Authors:** Gabriel Silva Santos, Roberto Salguero-Gómez, André Tavares Corrêa Dias, Maja Kajin

## Abstract

Until recently, natural selection was assumed to reduce temporal fluctuation in vital rates due to its negative effects on population dynamics – the so-called Demographic Buffering Hypothesis (DBH). After several failures to support the DBH in the two decades since it was first posited, an alternative hypothesis was suggested; the Demographic Lability Hypothesis (DLH), where population vital rates should track rather than buffer the environmental conditions. Despite the huge contribution of both hypotheses to comprehend the demographic strategies to cope the environmental stochasticity, it remains unclear if they represent two competing patterns or the extreme ends of a continuum encompassing all demographic strategies. To solve this historical debate, we unify several methods with an integrative theoretical approach where: i) using the sum of stochastic elasticity with respect to mean and variance – a first-order derivative approach – we rank species on a Buffering-Lability (DB-DL) continuum and ii) using the second-order derivative, we examine how vital rates are shaped by natural selection. Our framework, applied to 40 populations of 34 mammals, successfully placed the species on the DB-DL continuum. We could also link the species' position on the DB-DL continuum to their generation time and time to recovery. Moreover, the second-order derivative unveiled that vital rates with lower temporal variation are not necessarily under a strong pressure of stabilizing selection, as predicted by DBH and DLH. Our framework provides an important step towards unifying the different perspectives of DBH and DLH with key evolutionary concepts.

## Introduction

Modern demographic theory offers a wide array of hypotheses to explain the evolutionary processes that shape the diversity of demographic responses to environmental stochasticity (Charlesworth 1994, Pfister 1998, Healy *et al.* 2019, Hilde *et al.* 2020). Of these, two stand out as the most promising in linking evolutionary processes to demographic patterns: the Demographic Buffering Hypothesis (DBH) and its alternative hypothesis, the Demographic Lability Hypothesis (DLH) (Koons *et al.* 2009, Jongejans *et al.* 2010, McDonald *et al.* 2017, Hilde *et al.* 2020). Together, the DBH and DLH (hereafter *DB-DL perspective*), represent two contrasting ways by which natural populations cope with environmental stochasticity by either buffering (DBH) or tracking (DLH) changes in environmental conditions (Morris *et al.* 2008, McDonald *et al.* 2017, Hilde *et al.* 2020).

Coined over two decades ago, the DBH postulates that vital rates (*e.g*. survival, fecundity) that have the greatest influence on population growth should buffer the negative effects of the environmental stochasticity by presenting lower temporal variation than less influential vital rates (Pfister 1998). Because local extinction risk typically increases with increasing fluctuations in population size (Ovaskainen and Meerson 2010), demographic buffering is hypothesized to be a result of natural selection. This evolutionary process reducing variation in vital rates, called environmental canalization (Debat and David 2001, Gaillard and Yoccoz 2003), is expected to be more intense in vital rates that contribute the most to population growth rate (Pfister 1998).

Demographic buffering is expected to be particularly important for species that are less capable to recover after perturbations, which have also a higher risk of extinction due to demographic stochasticity (*e.g*. long-lived trees and primates, Morris *et al.* 2008, 2011, Hilde *et al.* 2020). Contrastingly, the DLH postulates that some species might take advantage if their vital rates were labile enough to permit population size track rather than buffering the environmental stochasticity (Koons *et al.* 2009, 2014, Jongejans *et al.* 2010, Hilde *et al.* 2020).

The DB-DL perspective simultaneously reflects *demographic patterns, i.e.* how populations respond to the environmental stochasticity (Morris *et al.* 2008, Jongejans *et al.* 2010, McDonald *et al.* 2017), and *evolutionary processes, i.e.* how the life-histories of the individuals that make up those populations were shaped by natural selection (Pfister 1998, Gaillard and Yoccoz 2003, Koons *et al.* 2009). Both demographic and evolutionary facets are important to predict the eco-evolutionary dynamic in response to climate changes (Metcalf and Pavard 2007, Burns *et al.* 2010, Salguero-Gómez *et al.* 2018). Despite the importance of understanding demographic responses to environmental changes, existing approaches to assess the DBH and DLH present several constraints and can therefore hinder our understanding of the eco-evolutionary processes behind the DB-DL perspective (Box 1, 2).

After two decades of consolidating the methodological bases of the DB-DL perspective (Hilde *et al.* 2020), a lack of consensus on how to identify evidence in favor of the DBH and DLH still persists (Box 1). Moreover, the bases of the DBH and DLH have been mainly developed in terms of mathematical models rather than trait-based mechanisms (Hilde *et al.* 2020). Indeed, only few authors have suggested trait-based mechanisms to support the DB-DL perspective (Morris *et al.* 2008, McDonald *et al.* 2017). Of those, even the prevalence of DBH in long-lived species is not well supported (Morris *et al.* 2011, Reed and Slade 2012, McDonald *et al.* 2017). Additionally, other attempts have found no correlation between the DBH and DLH and other life-history traits (*e.g.* generation time), environmental variables, nor phylogenetic relationships (McDonald *et al.* 2017).

In this study, we show that the lack of patterns in the assessment of DBH and DLH arises from different interpretations of what the two main approaches to assess them consider as evidence for DBH and DLH. Here we will refer to these approaches as multi-species comparative approaches (*e.g*. Morris *et al.* 2008, McDonald *et al.* 2017) and (very often single population) hypothesis testing approaches (*e.g.* Reed and Slade 2012, Jäkäläniemi *et al.* 2013). We argue that these approaches represent two perspectives with very distinctive eco-evolutionary implications that should be disentangled to better understand the performance and the evolutionary processes of the buffering and lability demographic strategies. To better assess the multiple facets of the DB-DL perspective, we reconcile different methods developed over the last two decades in a unified framework. Finally, we demonstrate the insights provided by our framework to better understand how species and populations both respond to and are shaped by environmental stochasticity.

### Box 1. Standard protocol to assess the DBH and DLH

The usage of simple correlation analyses (particularly the Spearman’s correlation) has been the most frequent approach to assess the DBH and DLH ever since they were posited (Pfister 1998, Hilde *et al.* 2020). Yet, two competing interpretations exist regarding the outcome of such correlation analyses (below). The lack of agreement between the two interpretations highlights the lack of consensus on how the demographic buffering vs. lability strategies manifest themselves in life histories.

The two competing interpretations of the correlation analyses to assess the DBH and DLH are based on different perspectives. Some researchers are interested in unveiling the patterns and distribution of the two demographic strategies - DBH and DLH across the tree of life in a multi-species comparative approaches (Jongejans *et al.* 2010, McDonald *et al.* 2017). Contrastingly, other researchers who assess DBH and DLH in terms of hypothesis testing are interested in finding support for one of these hypotheses by examining how natural selection have shaped the variance in vital rates (Reed and Slade 2012, Rotella *et al.* 2012). While the hypothesis testing approach looks for evidence to corroborate the patterns expected by either DBH or DLH (*e.g.* Jäkäläniemi *et al.* 2013), the multi-species comparative approaches assumes that all demographic strategies should lie somewhere between completely buffered (DBH) and completely labile (DLH)(*e.g.* McDonald *et al.* 2017). However, despite the well documented evidence in favor of the DBH, only scarce evidence exists to support the DLH (Hilde *et al.* 2020). Furthermore, the quantitative assessment of DBH and DLH reflects these different views of the comparative and hypothesis testing approaches as we show below.

Consider the example in Fig 1, where a negative Spearman’s correlation coefficient of −0.45 with P > 0.05 is shown. Despite the non-significant results, patterns like this have been used as evidence of demographic buffering in the multi-species comparative approaches (*e.g*. Doak *et al.* 2005, McDonald *et al.* 2017). Thus, the correlation coefficient is used as an index to place populations into the continuum, ranging from completely demographically buffered (Spearman’s correlation *ρ* = −1) to completely demographically labile (*ρ* = 1), regardless of the P-value (McDonald *et al.* 2017) - hereafter, *Demographic buffering-lability continuum* (or *DB-DL continuum*). However, other authors interpret the absence of statistical support for the DBH as evidence in favor of the DLH (*e.g.* Jäkäläniemi *et al.* 2013) or the absence of any particular strategy, neither DBH nor DLH (*e.g.* Reed and Slade 2012).

**Figure 1.**
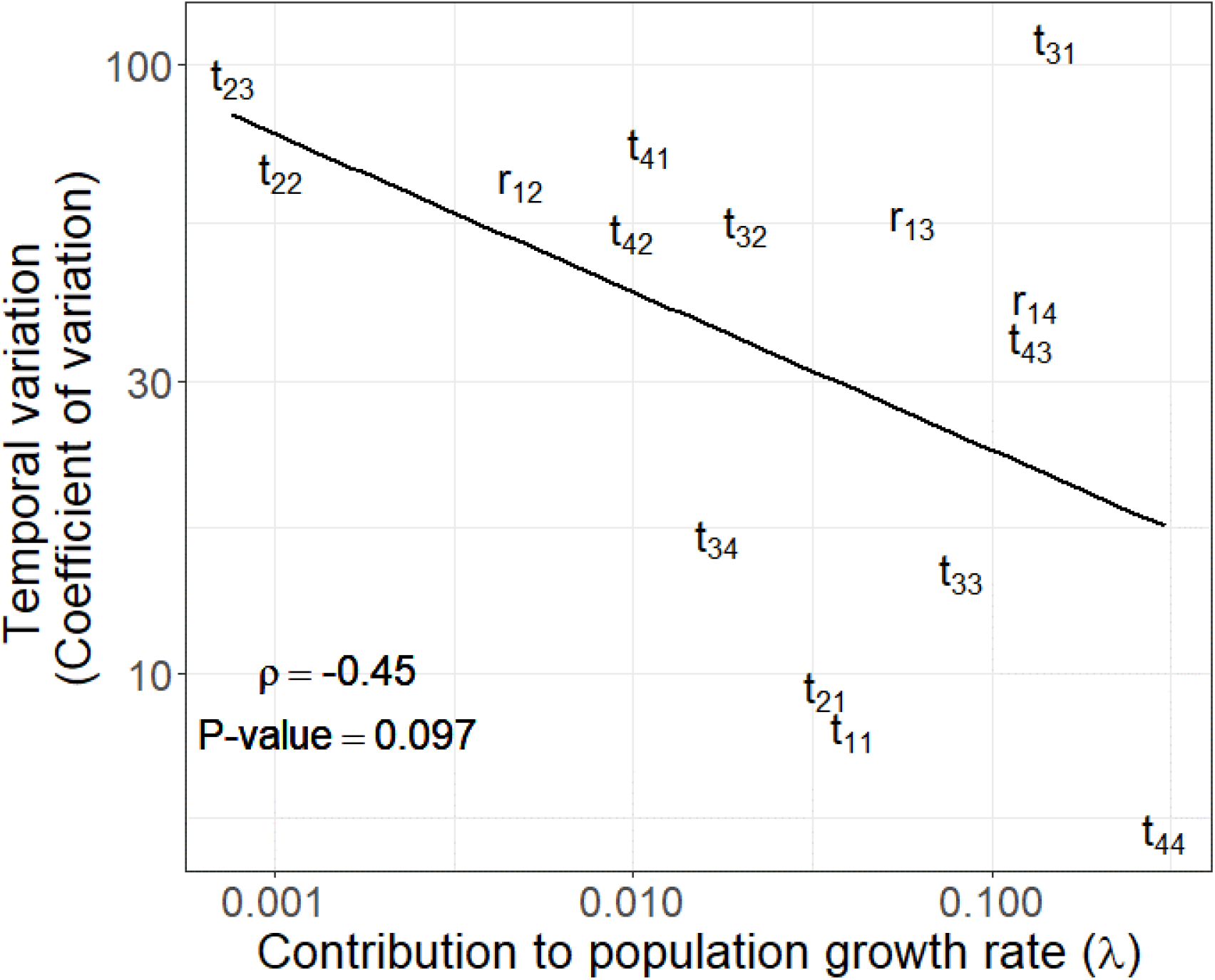
A hypothetical Matrix Population Model comparing the temporal variation in each matrix element. Letters represent (st-)age transitions (growth, stasis, and shrinkage – t) and recruitment (r), and the subscripts in the matrix element *a_ij_* correspond to transitions or recruitment events from stage *j* (column in matrix ***A***) in time *t*, to state *i* (row in ***A***) in t+1. The solid line represents the linear trend calculated by the least squares regression. The Spearman’s correlation coefficient (*ρ*) between the two variables and its P-value are informed in the figure.

### Disentangling demographic patterns and evolutionary processes in the DB-DL perspective: buffering mechanisms vs. canalization of variance

All populations are buffered in someway (Hunter 2009). Thus, the major question is not *if* but *to what extent* the populations are dependent on environmental stochasticity. Some traits, like endothermy or migration, can help individuals to buffer (or avoid) the negative effects of environmental conditions (Fuller *et al.* 2010, Edic *et al.* 2020). On the another hand, traits like fixed litter size and long parental care may be buffered, once they remain constant independently of environmental conditions (Meiri *et al.* 2020, Santos *et al. in prep*). All these traits which increase the capacity of an individual to buffer the effects of environmental stochasticity may be considered as buffering mechanisms (Santos *et al. in prep*). Ultimately, the capacity of the species to buffer the environmental stochasticity should be proportional to the set of buffering mechanisms of these species - including those traits which limit the population to increase in the good years (*e.g.* bonanzas).

Some patterns are expected to arise from these multiple traits. For example, bigger, long-lived species should be more buffered than smaller, shorter-lived species (Morris *et al.* 2008). Yet, such patterns do not tell us how these buffering traits have been shaped by natural selection (*i.e.* what processes led to these patterns). To assess the processes which underly these patterns, we need to analyze the *force and direction of natural selection acting on them*. Thus, if the temporal variation in an important buffering trait (*e.g.* longevity, age of first reproduction) exerts a negative impact on individual performance, its temporal variation might be canalized - *i.e. stabilizing selection* (Debat and David 2001) - increasing the population’s capacity to buffer the environmental stochasticity. Otherwise, natural selection might promote variation in the expression of these traits if they improve the individual performance in a variable environment, *i.e. disruptive selection* (Pasztor *et al.* 2000, Koons *et al.* 2009, 2014).

### A unified framework to assess the multiple facets of the DBH and DLH

Here, we propose a framework that overcomes the limitations of previous approaches (Box 1, 2) by explicitly disentangling demographic patterns from the evolutionary processes shaping the different vital rates. The application of this framework requires three steps which include two additional steps to the most largely used approaches to quantify DBH *vs*. DLH: the correlative analyses (Box 1). The **step 1** in the framework consists of comparing to what extent the life histories are variable and rank the populations of multiple species according to whether they are buffered or environmental dependent (labile) (Fig. 2A). **Step 2** and **step 3** are focused on populations rather than species. The **step 2** is analogous to the correlative analyses. However, we limited this step to visual inspection of biplots representing the relation between the temporal variance of vital rates and their contribution to population growth rate for each population. The graphical inspection of biplots helps to make inferences about the forces of natural selection acting on each vital rate before the empirical assessment by step 3. Finally, **step 3** enables to assess how temporal variance in survival and reproduction are shaped by natural selection (Figs. 2 B, C).

**Figure 2.**
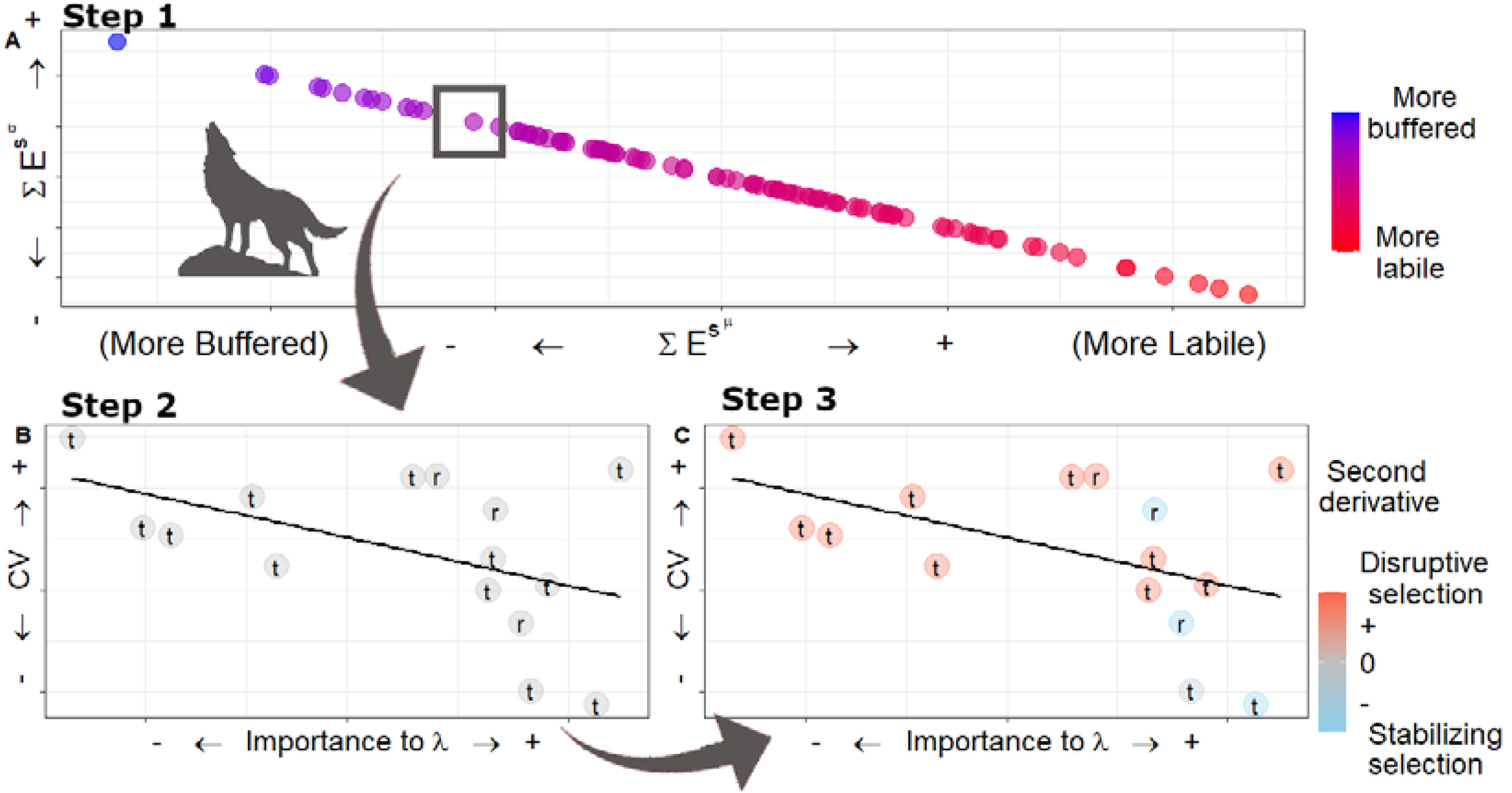
A unified framework placing life histories along a continuum of demographic buffering *vs.* demographic lability and linking the demographic processes to the underlying key evolutionary processes. Arrows show the flow between each step. Step 1 allocates (rank) each life history into the DB-DL continuum using the 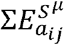 and 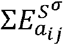. This step produces a diagonal line where species are placed according to their responses to environmental stochasticity, from the more buffered on the top left corner, to more labile on the bottom right corner. A population will move along the diagonal continuum, from the top left towards the bottom right corner, if its demographic processes *a_ij_* vary more across time – particularly the most contributive *a_ij_*. Dots represent 50 hypothetical life histories and colors represent the relative place along the continuum from more buffered (blue) to more labile (red). To perform steps 2 and 3, we must look at each population (each dot) in step 1, separately. Here we represent a hypothetical population from a hypothetical wolf species to assess whether the evolutionary process acting on its vital rates or demographic processes (or *a_ij_* for MPMs) correspond to expectations posit by the DBH and DLH. In step 2, we plot the elasticity of each vital rate or demographic process, *a_ij_*, *vs.* its coefficient of temporal variation providing a visual guide to evaluate the trend between temporal variance and its contribution to *λ*. Note that this step is common with most approaches that assess DBH and DLH, but, in our framework, it only serves to visually inspect the relationship. Finally, step 3 quantifies the type of natural selection driving the temporal variation in each demographic process, by obtaining the second-order derivatives, represented by different colors – from stabilizing selection (second-order derivatives < 0) in blue to disruptive selection (second-order derivatives > 0) in red.

Our unified framework provides a link between the DBH and DLH to evolutionary mechanisms, *i.e.* disruptive and stabilizing selection, where natural selection acts to reduce or increase the temporal variance in a specific vital rate (Fig. 2C). The full approach of our framework is explained in detail below, after a concise description of structured population models (in terms of both matrix population models [MPM hereafter; Caswell 2001] and integral projection models [IPM hereafter; Ellner *et al.* 2016]) - the common tools to assess the relative contribution of vital rates to population growth rate (de Kroon *et al.* 1986, Pfister 1998, Hilde *et al.* 2020).

#### The mathematical bases of the framework

The DBH and DLH have been mainly assessed by MPM (*e.g.* Pfister 1998, Rotella *et al.* 2012) and IPM (*e.g.* Rodríguez-Caro *et al.* 2020), which represent the analytical foundation for our framework. Both MPMs and IPMs are stage-structured, discrete-time demographic models (Caswell 2001, Ellner et al. 2016). In both, individuals in a population are categorized according to discrete (*e.g.* age, stage, as in MPMs; Caswell 2001) or continuous traits (*e.g.* size, height, as in IPMs; Easterling *et al.* 2000). For simplicity, for the rest of the framework description we focus on the usage of MPMs, but note that the same approaches are applicable to IPMs (*e.g.* Griffith 2017).

MPMs are matrix representations of life cycles (Ebert 1999). In an MPM **A**, each element *a_ij_* represents a demographic process, for example, a transition from a (st-)age to another or a contribution of a given (st-)age to the recruitment of new individuals. It is well known that each demographic process, *a_ij_*, contributes differently to the population growth rate (*λ*) depending on the life cycle and life history of the species (Silvertown *et al.* 1993). This contribution of *a_ij_* to *λ*, named *sensitivity*, can be assessed by simulating small changes on *a_ij_* or by the partial first-order derivative of each element in the matrix **A** with respect to *λ* (Caswell 2001, Tuljapurkar *et al.* 2003). As the sensitivities are dependent on the initial values of *a_ij_*, they are not comparable as some represent survival (bounded from 0-1) while others represent recruitment (≥0). Thus, to compare sensitivities for survival and recruitment, they must be rescaled to proportional contributions to *λ*, the so-called *elasticities e_ij_* (Caswell 2001).

As survival and recruitment are always, in some way, dependent on the current environmental conditions (Hunter 2009), we must expect (1) different values of *a_ij_* for each time *t* sampled and (2) a different population growth rate for each time *t*, *λ_t_* (Tuljapurkar *et al.* 2003). Moreover, in a stochastic environment, stochastic population growth rate (*λ_s_*) is a multiplicative process of bad and good years for the population (Morris and Doak 2002). Due to the multiplicative process that characterizes the *λ_s_*, the arithmetic mean of multiples 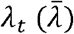 is not equivalent to *λ_s_* because the temporal dependence of *λ_s_* was not accounted for in 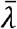 (Tuljapurkar *et al.* 2003). Indeed, very often 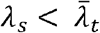 (Morris and Doak 2002). Additionally, elasticity in a stochastic perspective, namely stochastic elasticity, 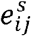 is composed of its mean and its variance, which both contribute to the stochastic population growth rate *λ_s_* (Tuljapurkar *et al.* 2003). Integrating these stochastic elasticities allow us to assess the effect of variation in each vital rate weighted by its relative contribution to *λ_s_* as explained below.

#### Comparing different life history responses to the environmental conditions

The sum of all stochastic elasticities 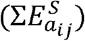 of an MPM **A** must be equal to 1 because it represents the contribution of all elements *a_ij_* to the stochastic population growth rate (*λ_s_*) (Tuljapurkar *et al.* 2003, Haridas and Tuljapurkar 2005). Moreover the sum of all stochastic elasticities 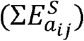 can be decomposed to assess how temporal variance and mean values of each vital rates contribute to *λ_s_* – they are respectively the *sum of stochastic elasticity with respect of the variance*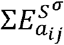, and the *sum of stochastic elasticity with respect of the mean* 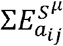 Once again, 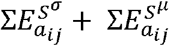 should be equal to 1 because 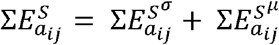.

The 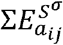, if applied to MPMs, might range from 0 to −1, producing a continuum of species’ responses that is similar to the predictions provided by the DBH and DLH (Box 1) – the *DB-DL* continuum (Haridas and Tuljapurkar 2005, Morris *et al.* 2008). A population with 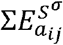 close to 0 - the *Demographic Buffered-end* (or *DB-end*) *-* has only a minor temporal variation in its vital rates, particularly in those vital rates that have a major contribution to *λ_s_*. The higher the variance in the most important vital rates, the further the population will be from 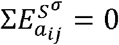(Fig. 2A). In other words, the further a population is from 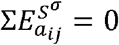, the closer it is to the *DL-end* of the DB-DL continuum. As the 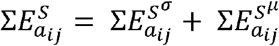, any change in 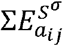 will reflect in 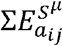, thus, both values will provide the same information.

The values of 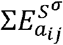 and 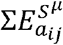 represent how species responded to the environmental stochasticity during the time sampled (Tuljapurkar *et al.* 2003). These values should vary from population to population according to the environmental conditions which populations were exposed to at the time, thus, the 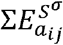 and 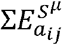 have also been referred as *the relative effect of environmental variation to population growth rate* (Morris *et al.* 2008). Yet, the 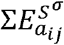 and 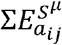 should vary according to species-specific range, because there are limits to how the most important vital rates can vary without the population becoming extinct, particularly regarding survival (*e.g.* Rodríguez-Caro *et al.* 2020). We also expect limits on reproductive output imposed by physiology and morphology of some species (*e.g.* Meiri *et al.* 2020). Ultimately, both 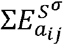 and 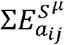 and the coefficient of correlation, can be used to place populations in a DB-DL continuum (Box 1). However, unlike previous methods, the 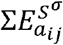 and 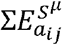 overcomes the key limitations characteristic of the correlative analyses and allows assessing DBH and DLH evidence more clearly (Box 2).

#### Revealing the evolutionary processes that shape the life histories

Evolutionary processes have been immediately evoked to explain the patterns arising from the comparative approach when assessing the position of species along the DB-DL continuum (Morris *et al.* 2011, McDonald *et al.* 2017). Thus, species placed at the DB-end of the continuum are commonly expected to be under strong selection pressures to reduce temporal variation in their most important vital rates (Gaillard and Yoccoz 2003). Contrastingly, natural selection is claimed to promote temporal variation in the most important vital rates in those species placed at DL-end of the continuum (Koons *et al.* 2009, 2014). Such evolutionary claims behind the DB-DL continuum are key to bridge the gap between demographic and evolutionary processes but remain unsupported by previous studies which failed to find consistent patterns (McDonald *et al.* 2017). To test these predictions, we use the evolutionary assumptions of stage-structured population models and their mathematical proprieties, as detailed below. Again, we focus on MPMs, but the same properties also emerge from IPMs (Ellner *et al.* 2016).

Matrix Population Models have been used numerous times to identify the evolutionary processes at the population level (*e.g.* Lande 1982, Coulson *et al.* 2006, de Vries *et al.* 2020). Because the mean values of the matrix elements *a_ij_* represent the mean performance of individuals at each (st-)age in surviving and transmitting their genes to the next generation, the population growth rate assessed by the MPM (*λ*) is assumed to be a proxy for the mean fitness of a population (Lande 1982, Brodie et al. 1995, Coulson et al. 2006, Metcalf and Pavard 2007, Engen et al. 2011). If *λ* can be used as a proxy for the mean fitness, each *a_ij_* is one of the fitness components and its contribution to the mean fitness can be quantified with its respective *sensitivity*. Nevertheless, the sensitivity obtained as the first-order derivative of *a_ij_*, assumes a linear relationship between a demographic process *a_ij_* and *λ*, even though this assumption is often violated in natural systems (Caswell 1996). The non-linearity between the contribution of each *a_ij_* and *λ* is adequately verified (and quantified) by using the second-order derivatives of pairs of *a_ij_* (Caswell 1996).

The assumption that *λ_s_* is approximately the long-term mean fitness in a population in a stochastic environment (*e.g.* Lande 1982, Coulson *et al.* 2006, de Vries *et al.* 2020), allows us to interpret the results from second-order derivative in terms of disruptive and stabilizing selection (Caswell 1996, McCarthy et al. 2008, Shyu and Caswell 2014). Therefore, if the second-order derivative of demographic process *a_ij_* is negative – representing a concave function (∩-shaped) - natural selection tends to reduce temporal variation in *a_ij_* (Brodie et al. 1995). We can expect a negative second-order derivative for demographic processes *a_ij_* with high contribution to fitness and low temporal variance, evidencing buffering as well as an environmental canalization process – a prediction from the DBH (Pfister 1998, Gaillard and Yoccoz 2003). On the other hand, if the second-order derivative is positive (U-shaped), the temporal variance of demographic process should increase, thus evidencing a disruptive selection, as predicted by the DLH (Pasztor *et al.* 2000, Koons *et al.* 2009, 2014).

### Box 2. Advantages of the unified analytical framework

Numerous limitations in using correlative analyses to assess the DBH and DLH have been pointed out since it was first proposed by Pfister (1998) (Morris and Doak 2004, Doak et al. 2005, McDonald et al. 2017). While some of these limitations have been overcome with corrective methods, others remain neglected. Below, we point out these limitations and how the framework proposed to handle them in contrast to the correlative analyses – the dominant approach to assess DBH and DLH (Table 1).

**Table 1.** Performance comparison between the framework proposed here and the largely used approaches to test DBH and DLH (correlative analyses).

|  | Largely used protocol | New framework proposal |  |
| --- | --- | --- | --- |
| Limitation \ Approach | correlative analyses | Sum of stochastic elasticity (step 1) | Second-order derivative (steps 2 and 3) |
| Mean-variance scale | Solved<sup>1,2,3,*</sup> | Accounted for | Not relevant |
| iid | Neglected<sup>4</sup> | Accounted for | Accounted for |
| Life history x vital rates | Accounted for but without consensus** | Accounted for, but each step of the present framework has a clear distinction between patterns and processes |  |
| Equal weight | Neglected | Accounted for | Not relevant |

\*There are multiple ways to solve the scale problem but there is no consensus of which one is the best ([<sup>1</sup>]Morris *et al.* 2011, [<sup>2</sup>]Bjørkvoll *et al.* 2016, [<sup>3</sup>]McDonald *et al.* 2017)
[<sup>4</sup>] Doak *et al.* 2005.
\*\* Two different approaches exist and are assessed using the same methods *i.e.* multi-species comparative approach and hypothesis testing approach (Box 1).

### Mean-variance scale

Different demographic processes cannot be compared because they have different magnitudes of mean and variance (Morris and Doak 2004). For instance, consider two survival rates 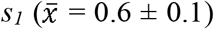 and (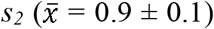. Both present similar variances, yet *s_1_* varies proportionally more than *s_2_*. Additionally, variation in *s_2_* is constrained by the maximum possible survival, which means that in some time intervals, no individuals died (*survival* = 1.0). A possible solution for such issues is rescaling variance by the maximum possible variance (Morris and Doak 2004, Morris *et al.* 2011). However, there is still no consensus on whether this is the best practice as different rescaling methods provide different results (Bjørkvoll *et al.* 2016, McDonald *et al.* 2017). Nevertheless, this is not a problem for our framework once vital rate variance is weighted by its proportional contribution to *λ* – the elasticity, a comparable metric between any kinds of demographic processes (Haridas and Tuljapurkar 2005).

### Independent and identically distributed (iid) variables

A common assumption in correlative analyses is the iid, which assumes that all demographic processes are independent from each other (Zar 2010). However, this might often not hold, since many vital rates can be linked by trade-offs (Doak et al. 2005). Thus, the selection pressure on a given demographic process (*e.g.* promoting variation) might affect other demographic processes in the opposite direction (Doak et al. 2005, Doak and Morris 2010). Since the data are not iid, the analytical framework must not be constrained by this assumption. In turn, our framework computes 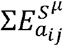 and 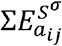, which inherently capture environmental and trade-off related variance. Additionally, the second-order derivatives (step 3), may also assess the force and direction of natural selection in trade-offs which is a potential source of correlation between different demographic processes (cross-correlative second-order derivative, not described here but see Caswell 1996, 2001, Shyu and Caswell 2014).

### Life histories *×* demographic processes or patterns *×* causality

Correlative analyses fail to provide a formal link between *processes* (or *causal mechanisms*) and *patterns* (Shipley 1999), especially because different demographic processes, which are under different selective pressures, are analyzed as part of the same pattern (Doak *et al.* 2005). For instance, Doak *et al.* (2005) demonstrate that even if high temporal variation in most of the demographic processes reduces *λ*, the variation of one or a few vital rates might have an opposite (and positive) contribution to *λ*. Mixing demographic processes under different evolutionary pressures in simple correlative analyses limits our understanding of the behavior of each demographic process and might lead to finding no pattern at all. Indeed, this might be a potential explanation of why most of the coefficients of correlation are close to zero in one of the most comprehensive uses of correlation analyses assessing DBH and DLH in plants by McDonald *et al.* (2017).

### Equal weight

Correlative analyses (Spearman’s correlation) are equally sensitive to demographic processes with high and low contributions to population growth rate. However, demographic processes with low contributions to population growth rate are free to vary under the predictions from the DBH and DLH (Doak *et al.* 2005). In this sense, it is preferable to think the predictions by DBH and DLH as *boundaries*, confining the maximum possible temporal variance of the demographic processes, rather than a trend which they should follow (Fig. 3). By using the step 1 of our framework, demographic processes with low contributions to *λ*, even if varying freely, should have minor contribution to place populations into the DB-DL continuum (Haridas and Tuljapurkar 2005). This characteristic of 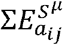 and 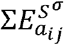fits conveniently with the predictions posted by the DBH and DLH.

**Figure 3.**
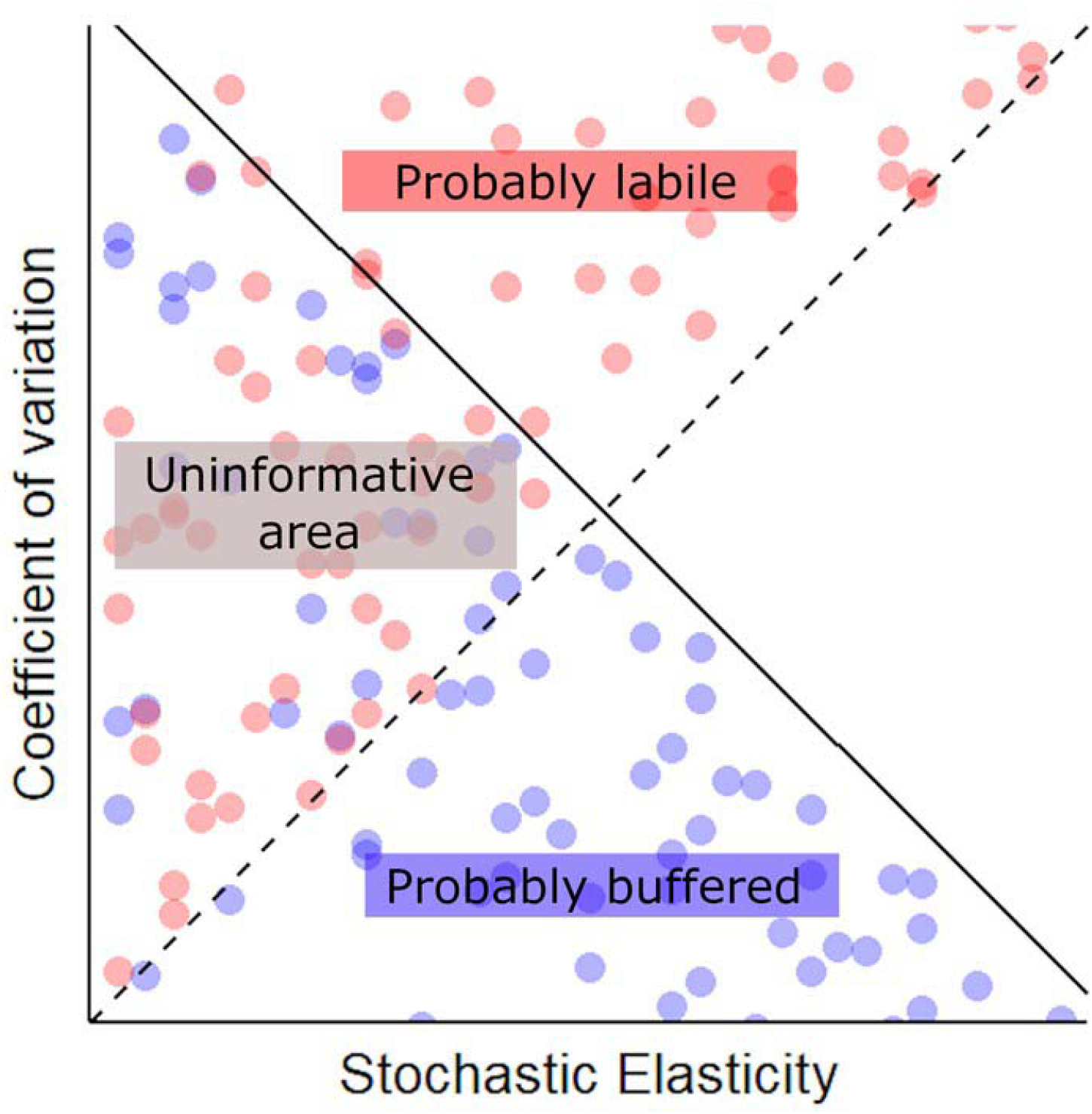
Theoretical relationship between stochastic elasticity and the coefficient of temporal variation under the predictions of DBH and DLH. The predictions from DBH and DLH are depicted in two different ways. Continuous and dashed lines represent a perfect support for the DBH and DLH, respectively, if we assume demographic processes should follow specific trends. Blue and red points represent the positions, where hypothetical vital rates should be placed, considering that natural selection acts by inducing boundaries to temporal variation under DBH and DLH predictions. Areas with co-occurrence of blue and red points together, cannot provide insights about the evolutionary process acting on demographic processe**s**, thus, they represent uninformative areas. Assuming the DBH and DLH as boundaries to temporal variation, the right quadrant is expected to be empty.

### Demographic buffering in mammals: a case study using the proposed framework

#### Study system and predictions

We demonstrate the performance of our framework to assess the DBH *vs.* DLH using MPMs from mammal species. Mammals are of special interest here for two reasons: first, mammalian life histories have been well studied (Gillespie 1977, Stearns 1983, Bielby *et al.* 2007, Jones 2011); and second, some of their populations have already been assessed in terms of DBH and DLH, particularly for primates (Morris *et al.* 2008, 2011, Reed and Slade 2012, Rotella *et al.* 2012, Campos *et al.* 2017). Together, the well-studied life histories and previous information about the DBH and DLH in mammals provide the necessary information to make accurate predictions to validate the performance of the proposed framework.

#### First step predictions – demographic responses in mammal life history

Long-lived species should be characterized by vital rates that are less dependent on environmental conditions (Gaillard and Yoccoz 2003, Morris *et al.* 2008, 2011). This might be particularly evident for primates, which have intense parental care, high longevities and flexible diets (Jones 2011). These aforementioned traits are expected to increase species’ capacity to buffer the negative effects of the environmental conditions. In addition, limited litter size imposed by physiological restrictions limits variation in recruitment even in the bonanza years (Campos *et al.* 2017). Therefore, we predict minimum variation on demographic processes for primates. The capacity to buffer the negative effects of the environmental stochasticity and physiological limitations should vary across the different lineages of mammals. Therefore, at the DL-end, we should expect short-lived species with a highly variable brood size, as is shown in small rodents (de Andreazzi *et al.* 2011) (Fig. 4).

**Figure 4.**
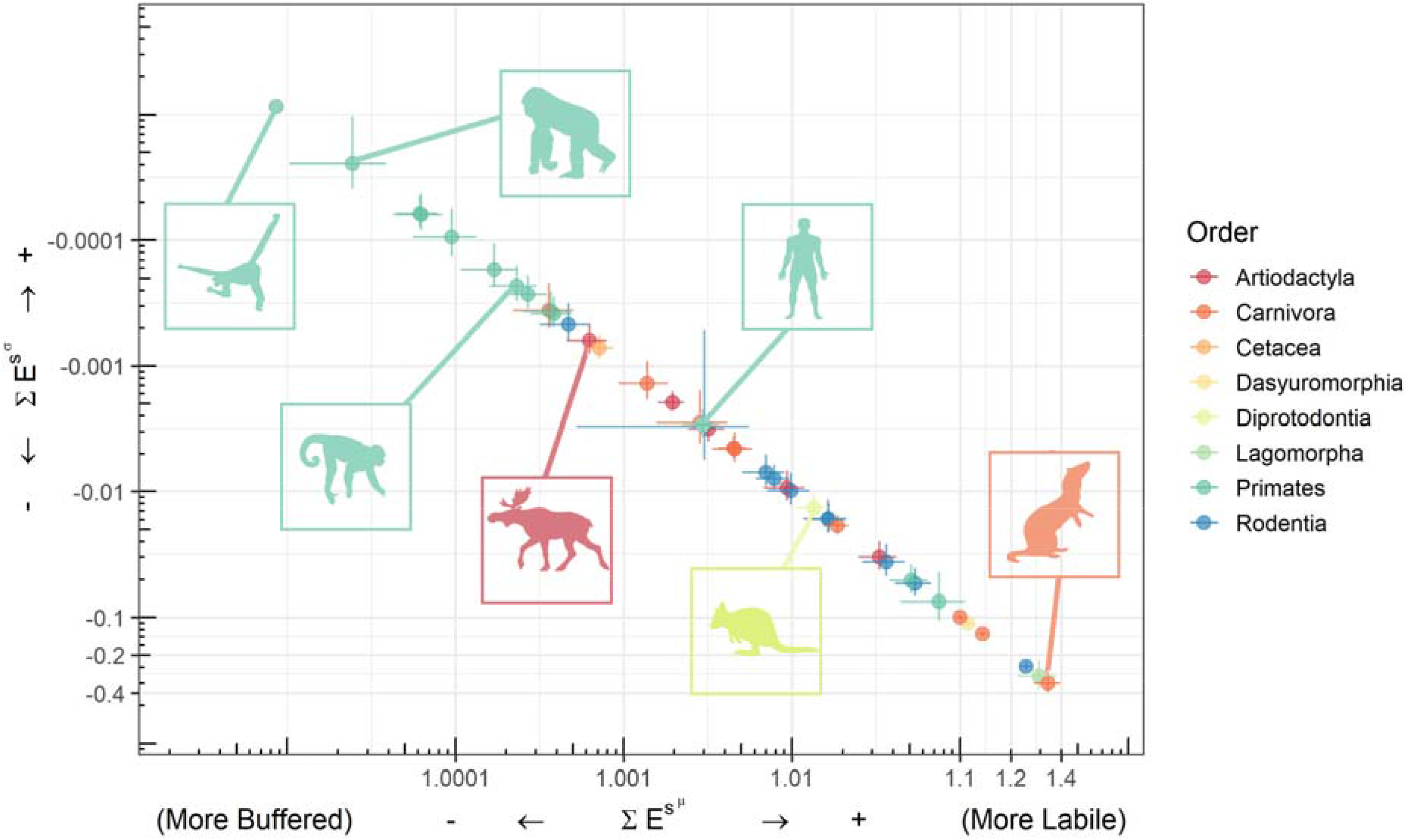
Results of the sum of stochastic elasticities with respect to their variance 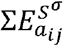 and mean 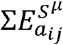 as suggested by our framework (step 1), ranking the 40 populations from 34 species of mammals from the COMADRE database onto a DB-DL continuum from more buffered (top left corner) to more labile (bottom right corner). Colors represent different taxonomic orders with Primates occupying the top left corner. Primate populations show the highest capacity to buffer environmental stochasticity, as expected.

Ultimately, we tested whether the demographic responses of mammals are related to their longevities and recovery capacities. The relationship between demographic responses has been suggested by Morris *et al.* (2008) but was not supported by McDonnald *et al.* (2017). On the other hand, a negative relationship between buffering and recovery rates has been suggested as an ecological reason why demographic buffer should evolve, yet this claim remains untested (Ovaskainen and Meerson 2010, McDonald *et al.* 2017).

#### Second and third step predictions – patterns of variance and natural selection in mamma’s life history

**Step 2:** As we posited in the predictions for step 1, it is unlikely that primates should be placed at the DL-end in the DB-DL continuum. Yet, previous studies have found no support for the idea that the most important vital rates vary less in primates – a pattern expected to support the DBH (Morris *et al.* 2011). Thus, we do not expect clear patterns from the relationship between temporal variation of the demographic processes and their contribution to *λ* (step 2), particularly for primates (Fig. 5). **Step 3**: This step is more critical in our framework because only humans (Caswell 1996) and a species of perennial herb (Shyu and Caswell 2014) had their demographic processes evaluated in terms of second-order derivatives so far. Yet, as the DBH and DLH predict, we should expect the most important demographic processes to be under stabilizing selection to reduce their temporal variation, supporting the DBH, or to be under disruptive section to increase temporal variation, supporting the DLH (Fig. 5).

**Figure 5.**
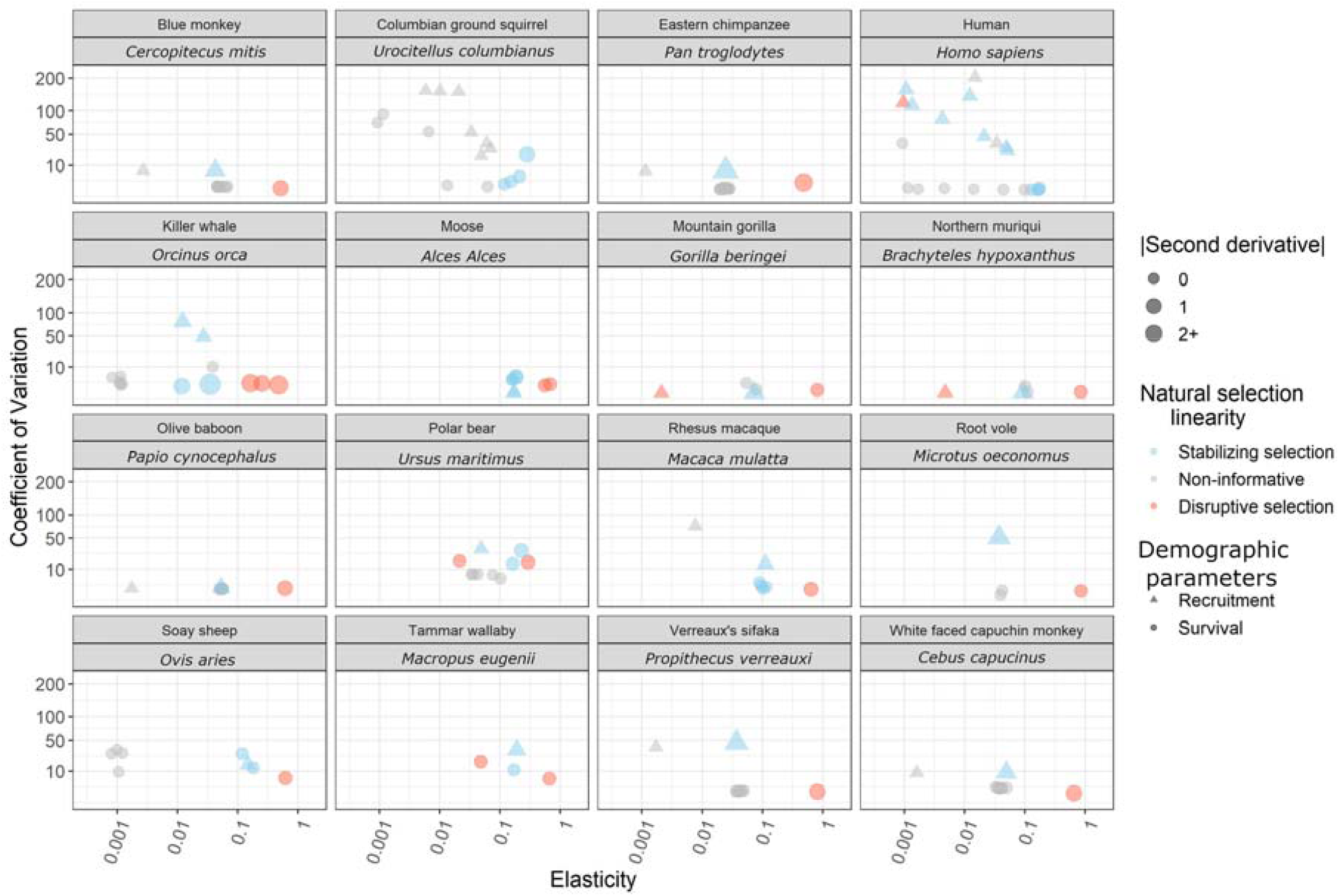
Results from steps 2 and 3 of the proposed framework (see Fig. 2B,C). The 16 populations presented here are those MPMs built by ages available in the COMADRE database. Different shapes represent different demographic processes (survival and recruitment). As in Figure 2B (step 2), the coefficient of temporal variation of each demographic process is plotted against its deterministic elasticity (see Box 1,2). We complete the graphical information with the value of the second-order derivative (denoted by size) and the direction of natural selection (denoted by color) acting on each demographic process represented by a specific matrix element. Full results are presented in supplementary material (Fig. S2).

#### Data and analyses

We used Matrix Population Models from 40 out of 139 studies with mammals available in the COMADRE database v.3.0.0 (Salguero-Gómez *et al.* 2016). These 40 populations encompass 34 species from eight taxonomic orders, and have been included in our analyses because they provide values of demographic processes (*a_ij_*) for three or more different moments, which allowed us to obtain the stochastic elasticity of each *a_ij_*. Although we are aware that not all possible temporal variation in vital rates may have been expressed within this time period, we assumed this was enough to provide sufficient variation for population comparison and to test and showcase our framework. Fortunately, several long-lived species, characterized by low variation in their vital rates, were studied for a long time (*e.g.* some primates have been studied for more than 20 years – Morris *et al.* 2011). We also removed the populations where either only survival or only reproduction rates were reported, because we could not calculate the stochastic growth rate for these populations. A detailed description of the analyzed data and their original sources is available in supplementary material (Table S1).

We included *Homo sapiens* in our analyses because it is the only mammal species, where second-order derivatives have been applied (Caswell 1996), thus making them ideal for species comparisons. The data for *Homo sapiens* were pooled from 26 modern populations from different cities to construct a variance. Note that in this case we are not working with true temporal variance but a spatiotemporal variance instead.

A subset of 16 populations, which had their MPMs organized by age (plus *Homo sapiens*), were used to run steps 2 and 3 of our framework. We selected these populations because the complexity of life histories is summarized only to two demographic processes: survival (t) and contribution to recruitment of new individuals (r), which is equivalent to or approximately equal to the mean reproductive output for each age, depending on the organization of the matrices (Ebert 1999). The advantage of using such matrices is that there are only two kinds of demographic processes included, survival and recruitment, instead of multiple transitions from one stage to another. We provide the results for all 40 populations in the supplementary material (Fig. S2). As the non-linear aspect of the contribution of vital rates to individual fitness is often neglected, we decided to provide an example to explore in detail how this nonlinear relationship appears in life histories in the DB-DL context (Fig. S3).

To perform the step 1 of our framework, the 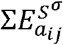 and 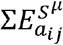, we followed Tuljapurkar *et al.* (2003). We checked if the position of species within the DB-DL continuum is correlated to longevity (here represented by generation time) and recovery rate (represented by damping ratio) using simple Spearman’s correlations. Generation time and damping ratio were extracted using the *Popbio* package (Stubben 2007). To perform step 2 of our framework, we calculated the coefficient of temporal variation of each demographic process corrected by its maximum variance, as suggested by Morris *et al.* (2011) (Box 2), and plotted the variance against the respective deterministic elasticity, also extracted using the *Popbio* package. All analyses were performed using R version 3.5.1 (R Core team, 2018).

Finally, to perform the step 3 of our framework, the second-order derivatives were adapted from *demogR* (Jones 2007) following Caswell (1996) and applied for the mean MPM. Most contributive demographic processes usually have higher absolute values of second-order derivative (Carslake *et al.* 2008) as they are expected to be under stronger pressure of natural selection (Brodie *et al.* 1995, Pfister 1998, Benton and Grant 1999). On another hand, second-order derivative rarely equals zero for the demographic processes, even for those less contributive demographic processes (Carslake *et al.* 2008, McCarthy *et al.* 2008, Shyu and Caswell 2014). However, less contributive demographic processes are either less important or more prone to have neglectable curvature (particularly because they are expected to be under less pressure of the natural selection [Pfister 1998]). As there is no approach to assess whether a second-order derivative is significantly different from zero or not (see McCarthy *et al.* 2008), we arbitrary rounded the values of second-order derivative for the first decimal place and considered zero as non-informative.

## Results

We ranked 40 populations from 34 species according to the cumulative impact of variation in demographic processes on *λ_s_* using the first step of our framework (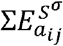 and 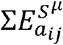). We show both values in Figure 4, but as the terms 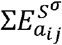 and 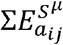are complementary and add up to 1, we only discuss the values from of 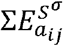for simplicity. Additional information is provided in the supplementary material (Table S1). The smallest contribution of variation in vital rates (*i.e.* maximum value of 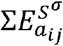, note that 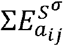 ranges from 0 to −1), meaning a less environmentally-dependent life history, was assigned to the Northern muriqui (*Brachyteles hyphoxantus*, Primates) (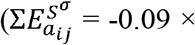 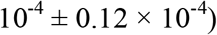) (mean ± standard deviation) and the most environmentally-dependent population was the stoat (*Mustela erminea*, Carnivora) (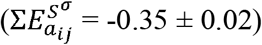).

All 14 primate populations were posited at the DB-end of demographic responses to environmental stochasticity, occupying the upper-left corner of the DB-DL continuum (Fig. 4). Other species in groups such as Carnivora (N = 10) and Artiodactyla (N = 5) are spread across the DB-DL continuum (Fig. 4). Our findings support that life histories with longer generation time are less affected by the environmental stochasticity, 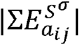 (Spearman’s correlation, *ρ* = −0.83, P < 0.01), which means, more buffered. On another hand, life histories more dependent of the environmental stochasticity have also higher recovery capacities, 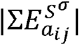 (Spearman’s correlation, *ρ* = 0.45, P = 0.01). These results are in concordance with the predictions from step 1 of our framework.

As we predicted for the step 2 (graphical inspection), we could not observe a clear negative correlation between the contribution to *λ* from separate demographic processes and their temporal variation. Particularly for primates, with the lack or minor temporal variation in demographic processes, an almost flat trend was created for most of the analyzed populations, *e.g.* Northen muriqui, and Montain gorilla (*Gorilla beringei*) (Fig. 5, S2). Yet, these abovementioned populations were all placed into the extreme DB-end using the first step of our framework (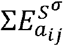 and 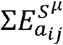). Such result highlights the importance of a clear definition of the expectations and the importance to choose the right method to assess the aspects of the demographic strategy in the wildlife populations. A full discussion is provided below.

By using the second-order derivative (step 3), our framework explicitly assessed the evolutionary forces of natural selection acting on each demographic process. However, no clear pattern arose from this step (Fig. 5, S2). Instead, we could identify some evidence of stabilizing or disruptive selection in almost one half of demographic processes (47% and 23 % respectively), mainly mixed within each life history (Fig S4). For instance, evidence of disruptive and stabilizing selection is present in the most important demographic parameters of White face capuchin monkey (*Cebus capucinus*), the rhesus monkey (*Macaca mulata*), and the tamar wallaby (*Macropus eugenii*) populations (Fig. 5, S4). We also use these populations to exemplify how variations in some of the most important demographic processes determine the long-term mean fitness (*λ_s_*) (Fig. S3). Only a few populations, like humans and Columbian ground squirrel (*Urocitellium colombianus*), presented negatives values of second-order derivatives for all most important demographic processes – a pattern expected by DBH (Fig. 3, 5). Similarly, only a few populations (*e.g.* killer whale [*Orcinus orca*]) presented all most important demographic processes under disruptive selection – evidencing DLH (Fig. 5, S4). A general profile on the frequency of what kind of natural selection the demographic processes are under is provided in the supplementary material (S4).

## Discussion

Our framework allows researchers to detect, quantify, and compare the extent to which natural populations are demographically buffered or labile in the environment, as well as the evolutionary processes shaping this capacity to buffer or track the environmental stochasticity (Fig. 2). In addition, our framework enables the development of an explicit link between the demographic buffering hypothesis (DBH; Pfister 1998) and the demographic labile hypothesis (DLH; Koons et al. 2009), and key evolutionary concepts such as disruptive and stabilizing selection (Fig. 2C). Below, we discuss the performance and contribution provided by each step of our framework to unveil the patterns and processes behind the DB-DL perspective.

### Framework performance

By using the sum of stochastic elasticity of the stochastic population growth rate with respect of the mean 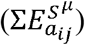 and variance (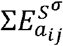, step 1), our framework supports the existing link between species’ buffering capacity, their capacity to recover after disturbances, and their longevity (Boyce *et al.* 2006, Morris *et al.* 2008, McDonald *et al.* 2017). Particularly for the recovery rate, our results represent the first empirical evidence between species’ recovery capacity and the DB-DL perspective. This particular result signifies an important gain, as both, the DBH and DLH, are mainly considered in terms of species’ capacity to track environmental conditions - particularly for the DL-end’s species (*e.g.* McDonald *et al.* 2017, Hilde *et al.* 2020). In terms of the relationship between longevity and the DB-DL continuum, our findings support recent work. Morris *et al.* (2008), also using 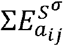 and 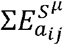, found that longevity determines a species’ position along the DB-DL continuum for 36 species of plants, mammals, and other vertebrates. Together, the emerging evidence is starting to indicate that life-history traits should drive the species’ position in the DB-DL continuum regardless of the taxonomic group. It is worth pointing out that other attempts, specifically those using correlative analyses (*e.g.* Morris *et al.* 2011, Reed and Slade 2012, McDonald *et al.* 2017)), did not reach the same conclusions.

The contrasting results of the correlative analyses versus the 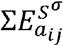 and 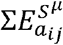, highlight the importance of carefully defining what should be expected for buffered or labile life histories. By clearly stating what we should expect based on the DBH and DLH, our framework using the 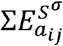 and 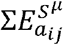(step 1 of our framework; Fig. 2), ranks species on a DB-DL continuum even when all demographic processes present low (or minimum) temporal variance. Temporal variance in vital rates is a necessary component to assess the DBH and DLH predictions and to rank species by using correlative analyses. Yet, several populations – particularly from highly buffered species – should present low (or minimal) temporal variance (*e.g.* Morris *et al.* 2011). A flat trend between temporal variation of vital rates and their contributions to *λ* might arise if all vital rates present similar and low temporal variation, which cannot be interpreted in terms of DBH and DLH using correlative analyses. Examples of flat or quasi-flat trends appear several times when visually representing the temporal variation of demographic processes and their contributions to *λ*. Such a pattern is not surprising for primates, where prolonged parental care and limited brood-size keep juvenile survival and adult reproductive rates constant in good or bad years (Morris *et al.* 2011; Campos *et al.* 2017, Salguero-gómez 2021).

Finally, the second-order derivative, step 3 in our framework (Fig. 2), reveals that the role of natural selection shaping temporal variation in vital rates is much more complex than expected by DBH and DLH. Indeed, we found that demographic processes within a population are often under a mix of stabilizing and disruptive selection - a pattern already suggested by Doak *et al.* (2005). Two findings here deserve further attention: firstly, stabilizing selection is not universal but is still dominant in the most important demographic processes. In fact, evidence for demographic processes under the influence of stabilizing selection might reach up to 46%, if we consider only those 25% of most contributing demographic processes in the life-histories. Such a result might help explain why evidence for the DBH has been dominant in the literature (see McDonald *et al.* 2017; Hilde *et al.* 2020). Secondly, we report evidence of beneficial variation - positive values of second-order derivative (U-shaped), even for most important vital rates with low temporal variation. Beneficial temporal variation in most important vital rates was suggested by Koons *et al.* (2009), particularly for short-lived organisms. Here, we show that natural selection might also increase the variance in demographic parameters in long-lived organisms. We found such pattern particularly for primates where most important demographic processes are the survival of oldest individuals (*e.g.* White face capuchin monkey [*C. capucinus*]). This result is probably a consequence of natural selection promoting longer lifespans in some populations, like those of primates, since we found positive second-order derivatives for adult survival of primates (Caswell 1996, Jones 2011, Jones and Tuljapurkar 2015, Colchero *et al.* 2016).

### General insights: challenges and future directions

Investigating what mechanisms drive species’ positioning along the DB-DL continuum is a promising avenue of further investigation. Our results from step 1, 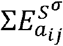 and 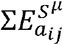, show that demographic buffering is a stronger pattern in Primates than Carnivora and Artiodactyla, for example. This pattern in some but not other taxa challenges the view that demographic strategies may not be predicted via phylogenetic distance as suggested by Morris *et al.* (2008). While we did not test this hypothesis explicitly here, however, our trait-based prediction to species’ buffering capacity supports the idea that more phylogenetically related species (or species with more similar traits) should present more similar buffering capacities. Indeed, the clustering of Primates but not Carnivora – a widespread taxa in terms of longevity and reproduction rate (van de Kerk *et al.* 2013) – supports the idea that similar species should be placed closer on the DB-DL continuum. We suggest further research should focus on exploring how the DB-DL continuum manifests across the Tree of Life and the role that morpho-physiological and life-history traits play in determining the demographic strategies of these species.

The insights provided by the second-order derivatives used in our framework (step 3) represent an important step to integrate ecological and evolutionary implications of selection acting on vital rate temporal variance. The contribution provided by the second-order derivatives was critical to link the evolutionary predictions posited by the DBH and DLH (Pfister 1998, Gaillard and Yoccoz 2003, Hilde *et al.* 2020). Moreover, the second-derivatives provide the necessary information when simple linear relationship fails to predict the effect of temporal variation (perturbation) of a given demographic processes to the mean fitness (Carslake *et al.* 2008, Shyu and Caswell 2014). We suggest the second-order derivatives to become more integrated into the ecological and evolutionary aspects of the demographic analyses with the attempt to link trait-based demographic models. For example, Integral Projection Models (IPMs; Easterling *et al.* 2000) usually use continuous traits (*e.g.* size) as a proxy to predict survival and fecundity (Ellner *et al.* 2016). As such, the same approach presented here might be adaptable to assess the evolutionary forces shaping the temporal variation of continuous traits used to build the IPMs.

Regardless to the potential of second-order derivatives in stochastic demography, their application is still rather limited (but see Caswell 1996, Carslake *et al.* 2008, Shyu and Caswell 2014, 2016), and has for the most part focused on methodological aspects (*e.g.* Caswell 1996, Shyu and Caswell 2014). However, despite the progress in the field since the implementation of the second-order derivatives for structured population models led by Caswell (1996), this method had not yet been applied interspecifically for comparative purposes. Thus, in order to make such comparison, we designed a specific profile of each life history based on the characteristics of the second-order derivative values to make them comparable. Despite the success of our approach to perform descriptive analyses, this approach offer limited statistical inference and remains a challenge for the future.

## Conclusions

Both, the Demographic Buffering Hypothesis (DBH) and the Demographic Lability Hypothesis (DLH) have become important pieces of an ecological puzzle to predict population responses to environmental stochasticity, climate change, and direct anthropogenic disturbances (Pfister 1998, Boyce *et al.* 2006, McDonald *et al.* 2017, Vázquez *et al.* 2017). Yet, the ecological and evolutionary facets of the DBH and DLH remain tangled in different perspectives that could not have been properly assessed by correlative analyses, as done to date. As a consequence, the literature exploring these two hypotheses is full of contradiction (as reviewed in Hilde et al. 2020). Even comparing small groups of closely related species (*e.g.* primates [Morris *et al.* 2011, Campos *et al.* 2017], rodents [Reed and Slade 2012], or herbs [Jäkäläniemi *et al.* 2013]) the results can be contradictory. The framework we propose here disentangles the ecological and evolutionary facets of population responses to stochastic environments, and provides the necessary tools to quantify and link them through two steps (sum of stochastic elasticity of the stochastic population growth rate with respect of the mean 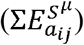 and variance 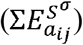 and the second-order derivative), in addition to the usual approach to assess the DBH and DLH (Pfister 1998, Morris and Doak 2004, McDonald et al. 2017).

Applied for 40 populations of different species, our framework helps to solidify classic and more recent avenues of research. Particularly, we highlight the capacity to link the DB-DL perspective with life-history traits (*e.g.* generation time), recovery rates and the possibility to link such results to the recent emphasis in linking functional traits and demographic performance (*e.g.* Adler *et al.* 2014, Salguero-Gómez *et al.* 2018). Thus, future investigations should benefit from this framework to explore, for example, how body mass and behavioral traits correlate with different demographic strategies in environmentally stochastic conditions. Furthermore, the usage of second-order derivatives in the DB-DL perspective represents a promising research avenue to explore the role natural selection to shape the temporal variation of vital rates in different taxa.

## Acknowledgements

This study was financed in part by the Coordenação de Aperfeiçoamento de Pessoal de Nível Superior - Brasil (CAPES) - Finance Code 001. We are grateful for Samuel Gascoigne for providing valuable comments to improve the science and English of the manuscript. RS-G was supported by a NERC Independent Research Fellowship (NE/M018458/1). MK thanks Prociência-UERJ (#363861) and Ad Futura – Public Scholarship, Development, Disability and Maintenance Fund of the Republic of Slovenia (#11013-16/2018) for their support.

## Appendix 1 Data available in COMADRE Version 2.0.1 and results from step 1

**Table S1.** present the metadata used in step 1 of our framework and the respective results presented in the paper. The first four columns represent the information from where Matrix Populations Models (MPMs) were extract exactly as presented in COMADRE 2.0.1. Column titles differ from the database as “SpeciesAuthorComadre” is equivalent to “SpeciesAuthor” and “SpeciesName” is equivalent to “SpeciesAccepted” in COMADRE 2.0.1. The remaining columns present the results of step 1, where we present the raw values of 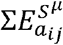 and 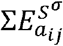, their respective standard deviation, population growth rate *λ*, number of available matrices (# matrices), generation time and damping ratio for each population. For ByAge, assign “TRUE” for MPMs built by age or “FALSE” if otherwise.

| SpeciesAuthorComadre | SpeciesName | CommonName | Order | $\Sigma E_{a_{ij}}^{S^{\mu}}$ | $\Sigma E_{a_{ij}}^{S^{\mu}}$ (sd) | $\Sigma E_{a_{ij}}^{S^{\sigma}}$ | $\Sigma E_{a_{ij}}^{S^{\sigma}}$ (sd) | # matrices | $\lambda$ | Generation time | Damping ratio | ByAge |
| --- | --- | --- | --- | --- | --- | --- | --- | --- | --- | --- | --- | --- |
| Homo_sapiens_subsp._sapiens | *Homo sapiens sapiens* | Human | Primates | 1.003 | 0.003 | 1.003 | 0.004 | 13.000 | 1.064 | 5.499 | NA | TRUE |
| Alces_alces | *Alces alces* | Moose | Artiodactyla | 1.001 | 0.001 | 1.001 | 0.001 | 13.000 | 1.205 | 10.628 | 2.168 | TRUE |
| Antechinus_agilis | *Antechinus agilis* | Agile antechinus | Dasyuromorphia | 1.111 | 0.111 | 1.111 | 0.011 | 2.000 | 0.931 | 2.153 | NA | FALSE |
| Brachyteles_hypoxanthus | *Brachyteles hypoxanthus* | Northern muriqui | Primates | 1.000 | 0.000 | 1.000 | 0.000 | 12.000 | 1.051 | 37.205 | 1.244 | TRUE |
| Callospermophilus_lateralis | *Callospermophilus lateralis* | Golden-mantled ground squirrel | Rodentia | 1.054 | 0.054 | 1.054 | 0.055 | 9.000 | 2.052 | NA | 3.955 | TRUE |
| Cebus_capucinus | *Cebus capucinus* | White faced capuchin monkey | Primates | 1.000 | 0.000 | 1.000 | 0.000 | 11.000 | 1.021 | 29.363 | 1.391 | TRUE |
| Cercopithecus_mitis | *Cercopithecus mitis* | Blue monkey | Primates | 1.000 | 0.000 | 1.000 | 0.000 | 14.000 | 1.036 | 21.452 | 1.240 | TRUE |
| Eumetopias_jubatus | *Eumetopias jubatus* | Northern sea lion; Steller sea lion | Carnivora | 1.005 | 0.005 | 1.005 | 0.002 | 2.000 | 0.904 | 9.645 | NA | FALSE |
| Felis_catus | *Felis catus* | Feral cat | Carnivora | 1.136 | 0.136 | 1.136 | 0.012 | 1.000 | 1.948 | 5.476 | 5.135 | FALSE |
| Gorilla_beringei | *Gorilla beringei* | Mountain gorilla | Primates | 1.000 | 0.000 | 1.000 | 0.000 | 21.000 | 1.027 | 38.817 | 1.239 | TRUE |
| Hippocamelus_bisulcus | *Hippocamelus bisulcus* | Huemul deer | Artiodactyla | 1.002 | 0.002 | 1.002 | 0.001 | 1.000 | 0.996 | 10.452 | 2.928 | FALSE |
| Lepus_americanus | *Lepus americanus* | Snowshoe hare | Lagomorpha | 1.294 | 0.294 | 1.294 | 0.165 | 2.000 | 0.812 | 2.290 | NA | FALSE |
| Lycaon_pictus | *Lycaon pictus* | African wild dog | Carnivora | 1.100 | 0.100 | 1.100 | 0.008 | 1.000 | 1.500 | 4.149 | 2.163 | FALSE |
| Macaca_mulatta_3 | *Macaca mulatta* | Rhesus macaque | Primates | 1.000 | 0.000 | 1.000 | 0.001 | 12.000 | 1.127 | 14.749 | 1.616 | TRUE |
| Macropus_eugenii | *Macropus eugenii* | Tammar wallaby | Diprotodontia | 1.013 | 0.013 | 1.013 | 0.012 | 7.000 | 0.981 | 4.872 | 10.930 | TRUE |
| Marmota_flaviventris_2 | *Marmota flaviventris* | Yellow-bellied marmot | Rodentia | 1.007 | 0.007 | 1.007 | 0.006 | 4.000 | 0.890 | 4.210 | 1.489 | FALSE |
| Marmota_flaviventris_3 | *Marmota flaviventris* | Yellow-bellied marmot | Rodentia | 1.008 | 0.008 | 1.008 | 0.005 | 4.000 | 0.921 | 5.017 | 3.575 | FALSE |
| Microtus_oeconomus | *Microtus oeconomus* | Root vole | Rodentia | 1.000 | 0.000 | 1.000 | 0.001 | 14.000 | 1.028 | 30.588 | 4.489 | TRUE |
| Mustela_erminea | *Mustela erminea* | Stoat | Carnivora | 1.334 | 0.334 | 1.334 | 0.117 | 2.000 | 1.258 | 5.803 | 11.201 | FALSE |
| Orcinus_orca_2 | *Orcinus orca* | Killer whale | Cetacea | 1.001 | 0.001 | 1.001 | 0.001 | 24.000 | 0.999 | 28.745 | NA | TRUE |
| Ovis_aries_2 | *Ovis aries* | Soay sheep | Artiodactyla | 1.033 | 0.033 | 1.033 | 0.020 | 3.000 | 1.099 | 8.036 | NA | TRUE |
| Pan_troglodytes_subsp._schweinfurthii | *Pan troglodytes* | Eastern chimpanzee | Primates | 1.000 | 0.000 | 1.000 | 0.001 | 22.000 | 0.982 | 37.652 | 1.132 | TRUE |
| Papio_cynocephalus | *Papio cynocephalus* | Olive baboon | Primates | 1.000 | 0.000 | 1.000 | 0.000 | 19.000 | 1.054 | 21.860 | 1.289 | TRUE |
| Peromyscus_maniculatus_2 | *Peromyscus maniculatus* | Deer mouse | Rodentia | 1.010 | 0.010 | 1.010 | 0.005 | 2.000 | 1.107 | 5.596 | 1.780 | FALSE |
| Phocarcos_hookeri | *Phocarcos hookeri* | New Zealand sea lion | Carnivora | 1.005 | 0.005 | 1.005 | 0.003 | 8.000 | 1.023 | 9.681 | 1.256 | FALSE |
| Propithecus_verreauxi | *Propithecus verreauxi* | Verreaux's sifaka | Primates | 1.000 | 0.000 | 1.000 | 0.000 | 12.000 | 0.986 | 21.963 | 1.376 | TRUE |
| Puma_concolor_8 | *Puma concolor* | Cougar | Carnivora | NA | NA | NA | NA | 10.000 | 1.115 | 4.806 | NA | FALSE |
| Rattus_fuscipes | *Rattus fuscipes* | Bush rat | Rodentia | 1.246 | 0.246 | 1.246 | 0.029 | 2.000 | 1.305 | 2.023 | NA | FALSE |
| Spermophilus_armatus | *Urocitellus armatus* | Uinta ground squirrel | Rodentia | 1.016 | 0.016 | 1.016 | 0.011 | 4.000 | 1.125 | 3.755 | 8.020 | FALSE |
| Spermophilus_armatus_2 | *Urocitellus armatus* | Uinta ground squirrel | Rodentia | 1.017 | 0.017 | 1.017 | 0.010 | 3.000 | 1.095 | 4.838 | 2.767 | FALSE |
| Spermophilus_columbianus | *Urocitellus columbianus* | Columbian ground squirrel | Rodentia | 1.036 | 0.036 | 1.036 | 0.025 | 3.000 | 1.009 | 4.508 | 2.510 | FALSE |
| Spermophilus_columbianus_3 | *Urocitellus columbianus* | Columbian ground squirrel | Rodentia | 1.003 | 0.003 | 1.003 | 0.006 | 3.000 | 1.200 | 4.524 | NA | TRUE |
| Ursus_americanus_subsp._floridanus | *Ursus americanus* | Florida black bear | Carnivora | 1.003 | 0.003 | 1.003 | 0.003 | 2.000 | 1.020 | 9.591 | 1.890 | FALSE |
| Ursus_arctos_subsp._horribilis_5 | *Ursus arctos* | Grizzly bear | Carnivora | 1.001 | 0.001 | 1.001 | 0.001 | 4.000 | 1.026 | 11.558 | 1.158 | FALSE |
| Ursus_maritimus_2 | *Ursus maritimus* | Polar bear | Carnivora | 1.019 | 0.019 | 1.019 | 0.007 | 2.000 | 0.941 | 18.491 | 1.552 | TRUE |
| Brachyteles_hypoxanthus_2 | *Brachyteles hypoxanthus* | Northern muriqui | Primates | 1.000 | 0.000 | 1.000 | 0.000 | 12.000 | 1.111 | 21.278 | 3.443 | TRUE |
| Cebus_capucinus_2 | *Cebus capucinus* | WhiteNAfaced capuchin monkey | Primates | 1.000 | 0.000 | 1.000 | 0.000 | 11.000 | 1.059 | 18.977 | 3.636 | TRUE |
| Chlorocebus_aethiops_2 | *Chlorocebus aethiops* | Vervet | Primates | 1.075 | 0.075 | 1.075 | 0.087 | 5.000 | 1.187 | 5.694 | 3.508 | FALSE |
| Erythrocebus_patas | *Erythrocebus patas* | Patas monkey | Primates | 1.051 | 0.051 | 1.051 | 0.038 | 5.000 | 1.128 | 3.499 | 2.291 | FALSE |
| Gorilla_beringei_subsp._beringei | *Gorilla beringei* | Mountain gorilla | Primates | 1.000 | 0.000 | 1.000 | 0.000 | 21.000 | 1.053 | 24.536 | 4.161 | TRUE |

## Appendix 2

**Figure S2.**
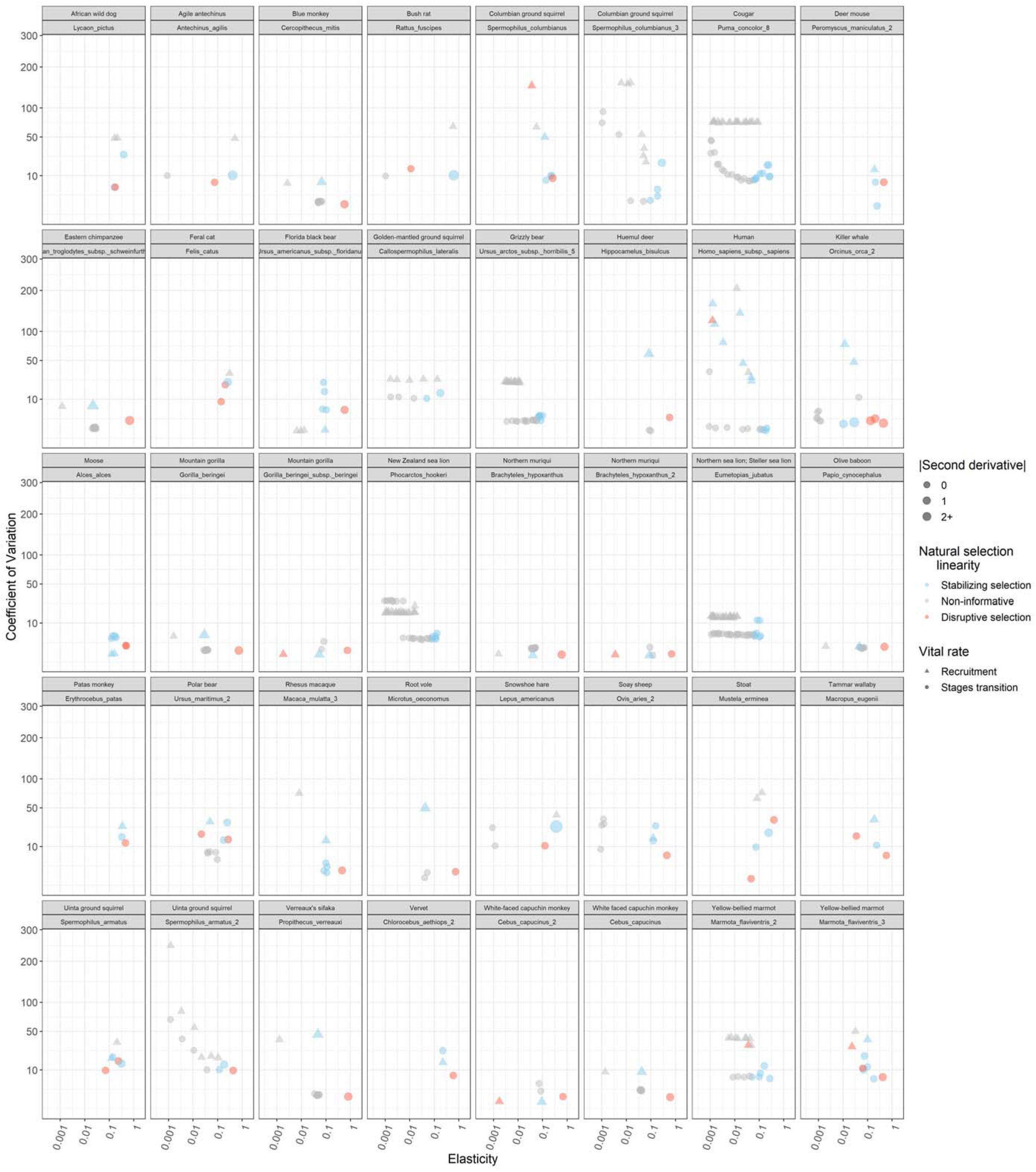
Results from steps 2 and 3 for all populations analyzed (see fig. 2B,C) in this study. Differently to figure 5, which only uses MPMs built by age with two kinds of transitions *a_ij_* – survival and recruitment, figure S2 presents MPMs built with all sorts of categories if they matched the criteria informed in *data and analyses* section. Therefore, instead of simple survival rates, we might disentangle it in other demographic processes as self-loops (retain in same stage) and jump for next stage (growth) are depicted in “stage transition”. Here we opted to simply inform what is a transition from a stage to another and what is contribution to recruitment. Moreover, differently to figure 5, common species names are paired with their respective identification in the “SpeciesAuthor” column in COMADRE database. Please note that the important results present in this figure have been covered by the discussion of results for figure 5.

## Appendix 3

**Figure S3.**
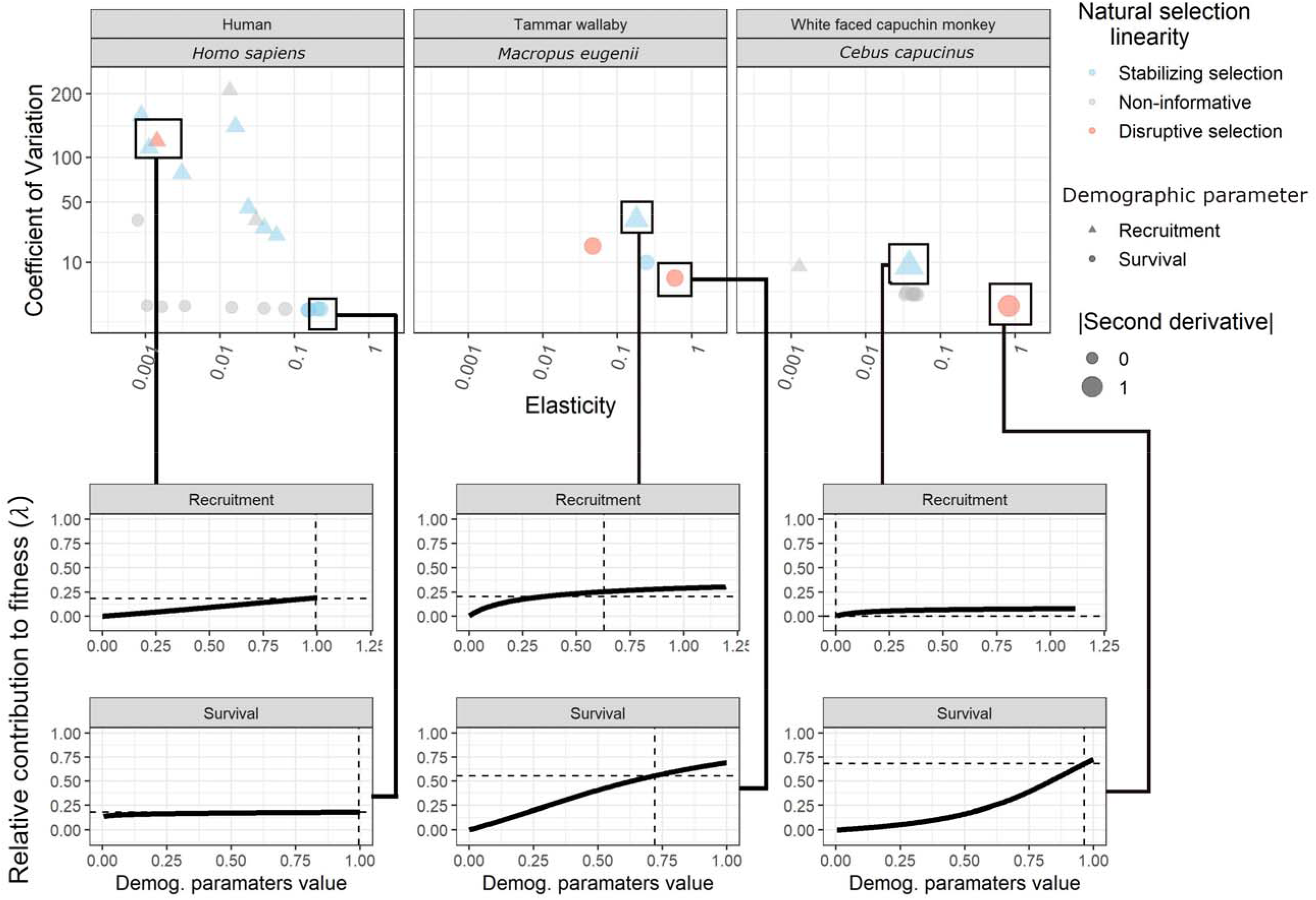
Example of nonlinear contribution to long-term mean fitness (*λ*) by two demographic processes in three populations analyzed in figure 5. The x-axis shows multiple simulated values of recruitment and survival with their respective contribution to long-term mean fitness (*λ*) on the y-axis. Vertical dashed lines show the recruitment and survival values from the mean MPM, as well as the corresponding value of *λ* (horizontal dashed line).

## Appendix 4

**Figure S4.**
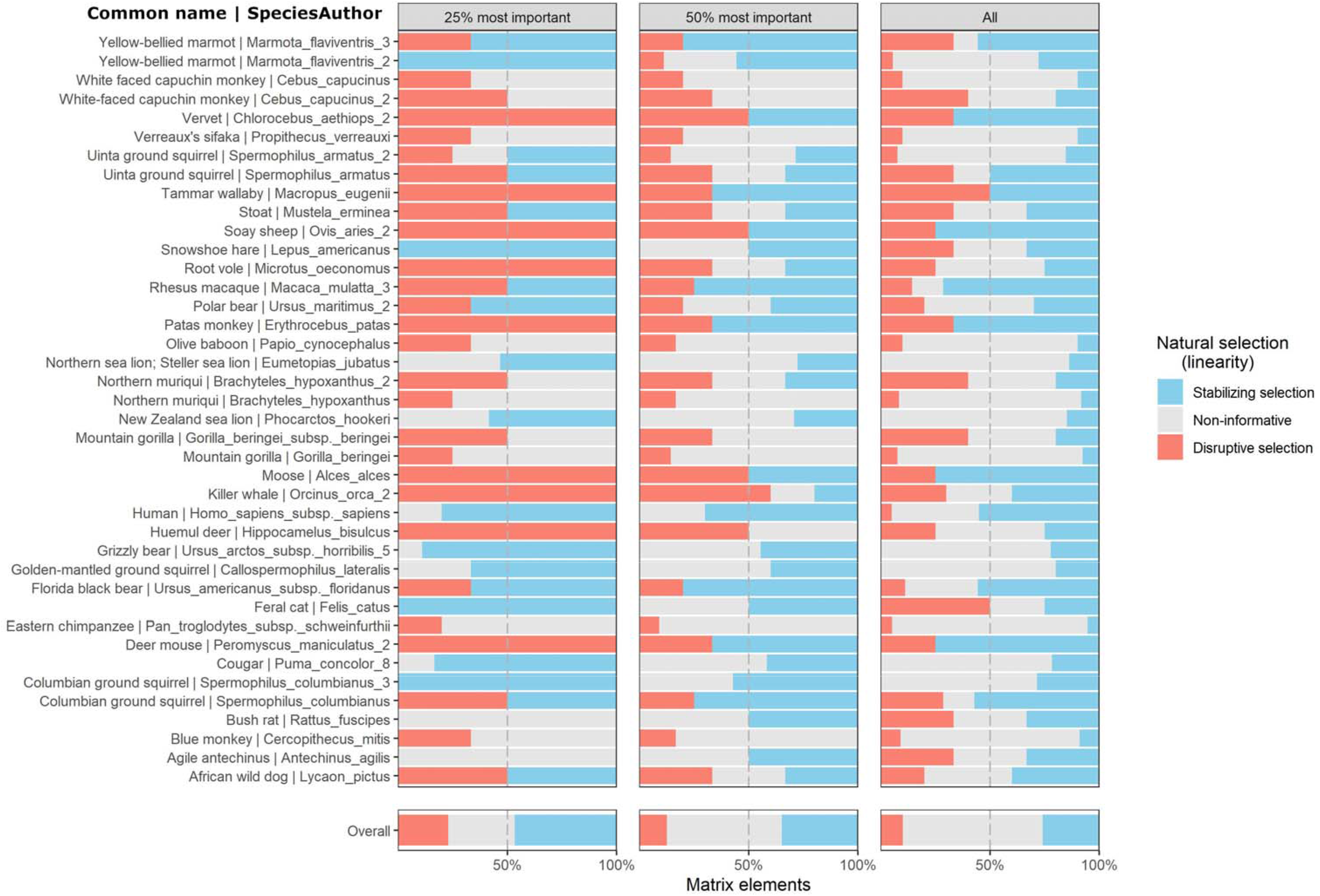
Most of life histories present mix of stabilizing and disruptive selection in their most important demographic processes and its proportion are presented for all life histories analyzed. From right to left, we present the proportion of disruptive and stabilizing selection acting on all demographic processes in a life-history (all non-zero entries in the MPM), followed by the proportion of each kind of natural selection for those 50% and 25% of most contributive demographic processes. At the end of each column the overall result including all life-histories is presented. Each life-history analyzed is informed at the left by species common name followed by its respective identification in the “SpeciesAuthor” column in COMADRE database.

